# Hierarchical syntax models of music predict theta power during music listening

**DOI:** 10.1101/2023.05.15.540878

**Authors:** Steffen A. Herff, Leonardo Bonetti, Gabriele Cecchetti, Peter Vuust, Morten L. Kringelbach, Martin A. Rohrmeier

## Abstract

Linguistic research showed that the depth of syntactic embedding is reflected in brain theta power. Here, we test whether this also extends to non-linguistic stimuli, specifically music. We used a hierarchical model of musical syntax to continuously quantify two types of expert-annotated harmonic dependencies throughout a piece of Western classical music: prolongation and preparation. Prolongations can roughly be understood as a musical analogue to linguistic coordination between constituents that share the same function (e.g., ‘pizza’ and ‘pasta’ in ‘I ate pizza and pasta’). Preparation refers to the dependency between two harmonies whereby the first implies a resolution towards the second (e.g., dominant towards tonic; similar to how the adjective implies the presence of a noun in ‘I like spicy…’). Source reconstructed MEG data of sixty-eight participants listening to the musical piece was then analysed. We used Bayesian Mixed Effects models to predict theta envelope in the brain, using the number of open prolongation and preparation dependencies as predictors whilst controlling for audio envelope. We observed that prolongation and preparation both carry independent and distinguishable predictive value for theta band fluctuation in key linguistic areas such as the Angular, Supramarginal, Superior Temporal and Heschl’s Gyri, or their right-lateralised homologues, with preparation showing additional predictive value for areas associated with the reward system and prediction. Musical expertise further mediated these effects in language-related brain areas. Results show that predictions of precisely formalised music-theoretical models are reflected in the brain activity of listeners.

## Introduction

Psycholinguistic studies overwhelmingly support that hierarchical syntactic structures, as formalised by recursive models of generative syntax, carry predictive value for behavioural as well as neurophysiological responses in humans during language comprehension (Bahlmann, Schubotz, & Friederici, 2008; Bornkessel, Zysset, Friederici, Von Cramon, & Schlesewsky, 2005; Brysbaert & Mitchell, 1996; Chen, Goucha, Männel, Friederici, & Zaccarella, 2021; Friederici, Friedrich, & Makuuchi, 2011; Meyer, Obleser, Anwander, & Friederici, 2012; Nelson et al., 2017; Nicol & Swinney, 1989; Zaccarella & Friederici, 2017). For example, a processing model for hierarchical syntax predicts increased working memory demand for pronouns with embedded antecedent (see Figure 1, bottom) compared to non-embedded antecedent (see Figure 1, top). Supporting this prediction, during EEG recordings, pronouns with embedded antecedents show systematically larger theta-power, a common neural marker of memory activity (Nyhus & Curran, 2010), compared to pronouns with non-embedded antecedents (Meyer, Grigutsch, Schmuck, Gaston, & Friederici, 2015).

**Figure 1.**
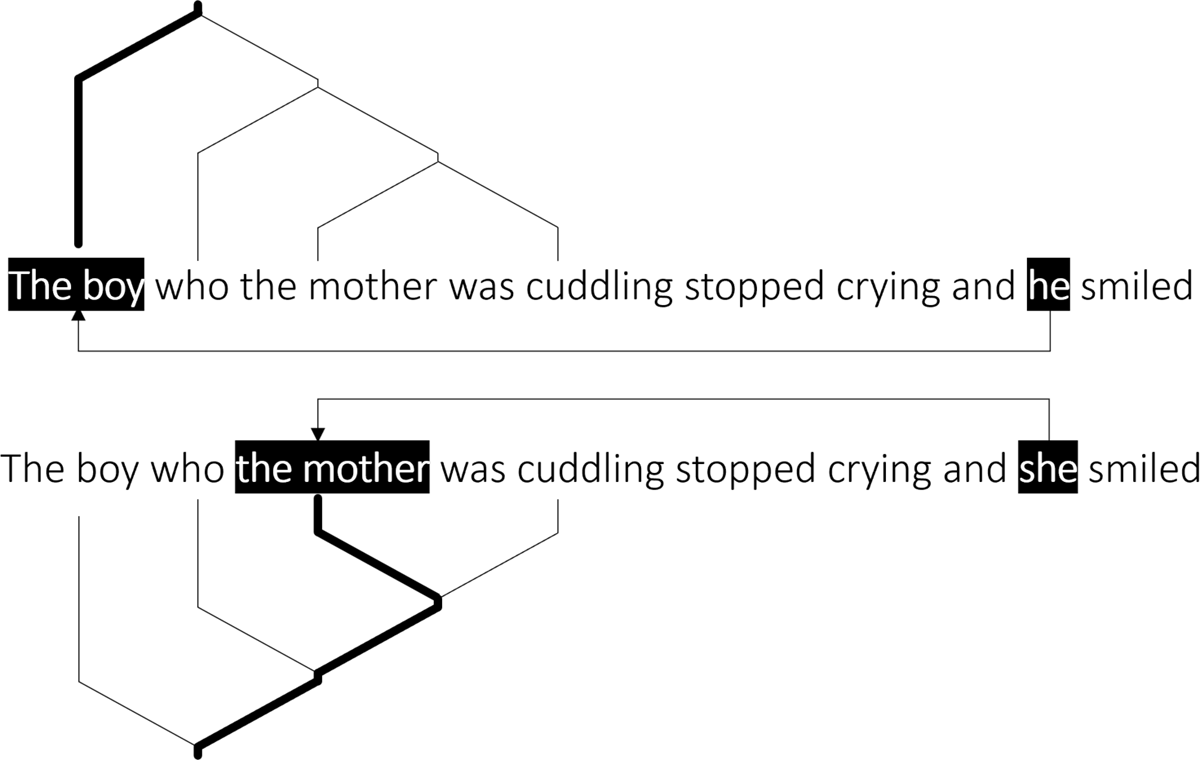
Example for a phrase with a non-embedded antecedent (top) and one with an embedded antecedent (bottom) after Meyer, Grigutsch, Schmuck, Gaston, & Friederici (2015). In the top sentence, the pronoun ‘he’ relates to the antecedent ‘the boy’, and ‘the boy’ is not grammatically embedded in the relative-clause constituent. In the bottom sentence, ‘she’ relates to ‘the mother’ which is embedded in the relative-clause constituent. Hierarchical syntax models predict higher working memory demand when processing pronouns with embedded antecedents compared to non-embedded antecedents.

However, hierarchical syntax models are not only adopted in the linguistic domain. Indeed, domains such as action sequences (Greenfield, Nelson, & Saltzman, 1972), dance (Charnavel, 2019), narrative (Van den Broek, 1988), or music (Keiler, 1978; Lerdahl & Jackendoff, 1983a) have also been described through the lenses of hierarchical syntactic models (for reviews, see Cohen, 2000; Fitch & Martins, 2014; Jackendoff, 2007). Yet, compared to the wealth of research in linguistics that shows the predictive and explanatory power of hierarchical syntax models for behaviour as well as neural activity, fewer studies have provided such evidence for non-linguistic stimuli. Here, we explore whether Meyer et al. (2015)’s findings – showing that hierarchical syntactic models systematically predict theta power as a function of embedding depth – also generalise to non-linguistic stimuli, specifically music. In the following, we first summarise some of the support for hierarchical syntactical structures in music, then we will review evidence supporting that theta-band activity is a promising neural marker for the predictions of hierarchical syntax models. We will then briefly describe the music-theoretically motivated formalism of musical hierarchical structures adopted here, before presenting novel results from the analysis of a MEG study, supporting the generalisation of Meyer et al. (2015)’s findings to the domain of music.

### Syntactic Structure in Music

#### Theoretical and formal models

The usefulness of modelling the experience of musical structure as computation (Cecchetti, Herff, Finkensiep, & Rohrmeier, 2020) to describe and explain structural relations in music has been explored from theoretical, computational, behavioural, and neuroscientific perspectives. From a theoretical perspective, describing musical structure in terms of hierarchical dependency relations in particular is a longstanding common practice (Lerdahl & Jackendoff, 1983b; Schenker, 1935). For example, theoretical accounts of Western harmony describe embedded structures, headed constituents, and nonlocal dependencies between chords (Aldwell, Schachter, & Cadwallader, 2018; Rohrmeier, 2011; Steedman, 1996) that would challenge purely local, non-hierarchical models – such as n-gram or Markov models without hidden states – due to the existence of embedded structures and long-term dependencies. Computational implementations of grammatical parsers further support the descriptive adequacy of hierarchical grammar models and show that harmonic sequences can be efficiently captured by hierarchical syntactic grammars (De Haas, 2012; De Haas, Rohrmeier, Veltkamp, & Wiering, 2009; Granroth-Wilding & Steedman, 2014; Harasim, Rohrmeier, & O’Donnell, 2018). Automatic grammar induction from musical corpora (Harasim, 2020; Harasim, O’Donnell, & Rohrmeier, 2021) further show that harmonic grammars can be learnt from musical pieces in an unsupervised fashion and approximate theoretical accounts and human annotations to a high degree (Harasim, Finkensiep, Ericson, O’Donnell, & Rohrmeier, 2020; Harasim et al., 2018; Herff, Harasim, Cecchetti, Finkensiep, & Rohrmeier, 2021). In summary, music theoretical accounts formalise some musical idioms, for example Western common-practice harmony (Lerdahl & Jackendoff, 1983a; Rohrmeier & Neuwirth, 2015), melodic lines (Finkensiep, Widdess, & Rohrmeier, 2019; Kirlin, 2014), polyphony (Finkensiep & Rohrmeier, 2021), rhythm (Rohrmeier, 2020b), or Jazz (Granroth-Wilding & Steedman, 2014; Rohrmeier, 2020a) through hierarchical syntactic models, and computational evidence supports the appropriateness of such accounts through direct implementation, unsupervised grammar inference, and corpus studies.

#### Behavioural evidence

From a behavioural perspective, listeners have been shown to perform structural revision, a feat commonly associated with linguistic garden-path sentences, supporting the notion that an abstract hierarchical representation of musical dependency structure exists in listeners (Cecchetti, Herff, & Rohrmeier, 2022). Musical phrases within a familiar idiom with clear syntactic interpretations under a generative model of tonal grammar also show a memory advantage (Cecchetti, Herff, & Rohrmeier, 2021), providing further support that the musical structures as identified by such syntactic models are predictive of listeners’ processing of musical phrases. Additionally, syntax models trained on expert annotations can predict behavioural responses when listeners are presented with interrupted musical phrases and asked to predict how many more chords are needed before the musical phrase is complete (Herff, Harasim, et al., 2021). This suggests that the hierarchical musical structures generated by recursive models of tonal syntax not only characterise the analytical understanding that musical experts have of musical pieces well, but also carry predictive value for human behaviour. Furthermore, the process involved in the processing of musical structure shares cognitive resources with the process involved in syntactic parsing in language (Fiveash & Pammer, 2014; Fiveash, Thompson, Badcock, & McArthur, 2018; Patel, 1998; Rohrmeier & Pearce, 2018; Van de Cavey & Hartsuiker, 2016), further highlighting the similarities between linguistic and musical processing of domain-specific structures. Taken together, this suggests that music theoretical accounts that describe music through hierarchical syntactic structure are not only useful when it comes to analysing music, but also allow predicting human behaviour by generating tangible hypothesis for crucial processing bottlenecks such as memory.

#### Neuroscientific evidence

Neuroscientific studies further support these behavioural findings. When comparing a version of a musical phrase that violates a syntactic nonlocal dependency with a version without such violation, an early negative brain response around 150ms (maximal around 220ms), as well as a later negative brain response around 500-850ms after chord onset emerges in EEG recordings (Koelsch, Rohrmeier, Torrecuso, & Jentschke, 2013). This provides an implicit neural marker that systematically varies with the violation of nonlocal dependencies as predicted by hierarchical syntactic models of music.

Furthermore, brain responses to functional harmonic dependencies are also subject to brain development during childhood. This was observed in a study showing that the aforementioned early negativity is already present in two-year old children whereas the later negativity only arises at a later developmental stage (Jentschke, Friederici, & Koelsch, 2014). Importantly, these neural markers also interact with the listeners’ familiarity with the musical system as well as with the complexity of the musical structure as captured by hierarchical syntax models. Specifically, more complex musical structures require higher levels of familiarity with the musical system before they can be detected using EEG (Ma, Ding, Tao, & Yang, 2018b). This further suggests that the neural activity of the marker is due to syntactic parsing of the musical structure, driven by a culturally acquired syntactic competence, rather than being purely driven by auditory processing regardless of musical structure. The interaction between structural complexity and listeners’ familiarity with a given musical idiom is particularly striking in neuroimaging studies of embedded musical structures. Ma, Ding, Tao, and Yang (2018a) showed that syntactic parsing is a learnable skill, and its acquisition coincides with changes in brain responses. Additionally, not only the presence or absence of long-distance dependencies shows a neural trace in EEG, but the kind of embedding also evokes noticeable differences. Specifically, the previously mentioned markers – that is, early and late negativities as well as late positive components (L. Sun, Feng, & Yang, 2020) – are sensitive to embeddings containing shifts to a musical key far away from the original tonal context compared to embeddings containing shifts to keys closer to the original context (Ma, Tao, & Yang, 2022). Taken together, these studies provide an encompassing view that support hierarchical syntactic accounts of musical structure as predictors of both behavioural as well as neurophysiological responses.

#### Brain regions involved in musical syntactic processing

In terms of brain regions of interest, Patel (2003) suggests that *‘linguistic and musical syntax share certain syntactic processes (instantiated in overlapping frontal brain areas) that apply over different domain-specific syntactic representations in posterior brain regions’* (p. 679). This notion is mirrored and generalised to other domains in a recent literature review by Friederici (2020). The conclusion is drawn from observations in patients with brain lesions that allow to clearly dissociate linguistic and musical syntax (Ayotte, Peretz, & Hyde, 2002; Ayotte, Peretz, Rousseau, Bard, & Bojanowski, 2000; Griffiths et al., 1997; Luria, Tsvetkova, & Futer, 1965; Peretz, 1993; Peretz et al., 1994; Sammler, 2009) combined with prior MEG and fMRI studies showing that both Broca’s (Maess, Koelsch, Gunter, & Friederici, 2001; Tillmann, Janata, & Bharucha, 2003) as well as Wernicke’s area (Koelsch, Gunter, et al., 2002), which are traditionally associated with linguistic processing, are also involved in the processing of musical harmony. A recent study further points towards the role of the inferior frontal gyrus – that contains Broca’s area—when it comes to processing of musical structure. Cheung, Meyer, Friederici, and Koelsch (2018) showed that the right inferior frontal gyrus in particular reflects processing of non-local dependencies by showing difference in fMRI activity when participants are listening to musical phrases that contain grammatical violations of a previous learned artificial grammar. In summary, key brain areas associated with syntactic parsing in language appear to also be involved in the processing of music (however also see X. Chen, et al. 2021; Chiappetta, Patel, & Thompson, 2022).

Whilst recent years have seen increasing behavioural as well as neuroscientific evidence for hierarchical syntactic parsing in music, these studies were predominantly concerned with showing proofs-of-concept for the existence of key constructs such as revision (Cecchetti, Herff, & Rohrmeier, 2022), sensitivity to violations of long-term dependencies (Koelsch, Rohrmeier, et al., 2013), or differentiation between embedded and non-embedded structures using short musical phrases (Ma et al., 2022). This was crucial to meet the necessary criteria to establish the appropriateness of hierarchical syntactic accounts of music. However, we argue that the support in the literature, summarised above, is now sufficient to warrant in-depth exploration of specific hierarchical accounts of music. This allows us to link concrete parameters of music-theoretical accounts to behavioural as well as neurophysiological responses to further our understanding of the perception and cognition of musical structure. Specifically, as music theoretical accounts allow to quantify embedding depths, and embedding depths predicts higher working memory load, we here aim to investigate whether depth of embedding in musical structure is predictive of theta power, a neural marker of memory activity, in MEG recordings.

### Working memory and Theta-band brain activity

#### Theta in language

As demands on working memory increase with the number of items that have to be distinguished, so does frontal-midline theta-band power (Gevins, Smith, McEvoy, & Yu, 1997). In general, theta-band power fluctuations have been associated with a wide variety of memory tasks, such as retention of letters (Krause et al., 2000), digits (Jensen & Tesche, 2002), words (Klimesch et al., 2001), and ordered spatial (Roberts, Hsieh, & Ranganath, 2013) and visual sequences (Hsieh, Ekstrom, & Ranganath, 2011; Liebe, Hoerzer, Logothetis, & Rainer, 2012). Theta power during encoding can also be used to predict subsequent memory performance (Khader, Jost, Ranganath, & Rosler, 2009; Sederberg, Kahana, Howard, Donner, & Madsen, 2003). The close association of theta activity with memory has been topic of several literature reviews, for example, by Hsieh and Ranganath (2014); Kahana, Seelig, and Madsen (2001); Klimesch (1999); Mitchell, McNaughton, Flanagan, and Kirk (2008); Nyhus and Curran (2010). As summarised in the previous section, an increase in open dependencies during parsing of syntactic structures is associated with rising demands on the memory system (Lewis, Vasishth, & Van Dyke, 2006). This makes theta activity a prime candidate to investigate hierarchical syntactic models, as such models make clear predictions about fluctuations in memory load. Indeed, in the linguistic domain this has already been explored extensively (for reviews, see Bastiaansen & Hagoort, 2006; Bastiaansen & Hagoort, 2010; Meyer, 2018).

A recent MEG study showed that theta activity systematically tags syntactic violations when participants read sentences (Pu, Cheyne, Sun, & Johnson, 2020). Theta power also reflects children’s development, with young children showing widely distributed theta activity during sentence comprehension, whereas sentence-comprehension-related theta power is predominantly localised in left central-posterior areas by adolescence (Maguire et al., 2022). Röhm, Klimesch, Haider, and Doppelmayr (2001) showed that syntactic aspects of sentence comprehension are reflected in the theta band, whereas semantic processing or processing of unstructured word lists is not (Bastiaansen et al., 2010). Theta activity has also been shown to be indicative of the high working-memory load during subject-object relative clauses (Weiss et al., 2005). This finding is particularly relevant to the present study, as it shows that theta power can be used as a marker of embedding depth during sentence processing. Taken together, these studies provide strong support for the use theta activity as a marker of syntactic processing.

#### Theta activity in music

In music, theta has been associated with reward processing and pleasantness (Ara & Marco-Pallarés, 2020, 2021; Sammler, Grigutsch, Fritz, & Koelsch, 2007), phrase boundaries (Silva, Barbosa, Marques-Teixeira, Petersson, & Castro, 2014), harmonic violations (Carrus, Koelsch, & Bhattacharya, 2011; Ruiz, Koelsch, & Bhattacharya, 2009), as well as audio envelope tracking (Miyazaki, Thompson, Fujioka, & Ross, 2013; Zuk, Teoh, & Lalor, 2020). However, its potential as a marker of embedding depths predicted by music-theoretical hierarchical syntactic models – to the best of our knowledge – has not yet been investigated. In order to test whether theta power reflects embedding depths, we need to adopt a formalism of musical structure that enables quantification of embedding depths. Here, we utilise the framework established in Rohrmeier and Neuwirth (2015) and Rohrmeier (2020b). We only briefly summarise this framework in the following, however, we describe in detail how it can be used to generate specific predictions for the embedding depths.

### Hierarchical Syntactic Model of Music in the Present Study

Hierarchical syntactic models of music are based on the understanding that musical events, such as notes or chords, may be recursively elaborated through the application of a limited set of generative rules (Baroni, Maguire, & Drabkin, 1983; Rohrmeier, 2011; Schenker, 1935). Structures arising through such recursive application of generative rules embody the dependencies between events, regardless of whether or not such events are adjacent in time. This is because the recursive application of rules allows for the possibility of embedded structures. The result of the recursive application of the rules can then be represented through tree analyses, where each branching corresponds to a rule application generating two children-nodes from a parent-node (see Figure 2 for a musical example). Vice versa, *parsing* a sentence or musical phrase can be understood as inferring the dependency structure, or parse-tree, of a given surface sequence, i.e., which chord dependencies (rule applications) may have generated the observed surface. Successful parsing depends on conscious or unconscious application of the grammatical rules that characterise a certain musical style, in analogy to the notion of syntactic competence (Chomsky, 1965).

**Figure 2.**
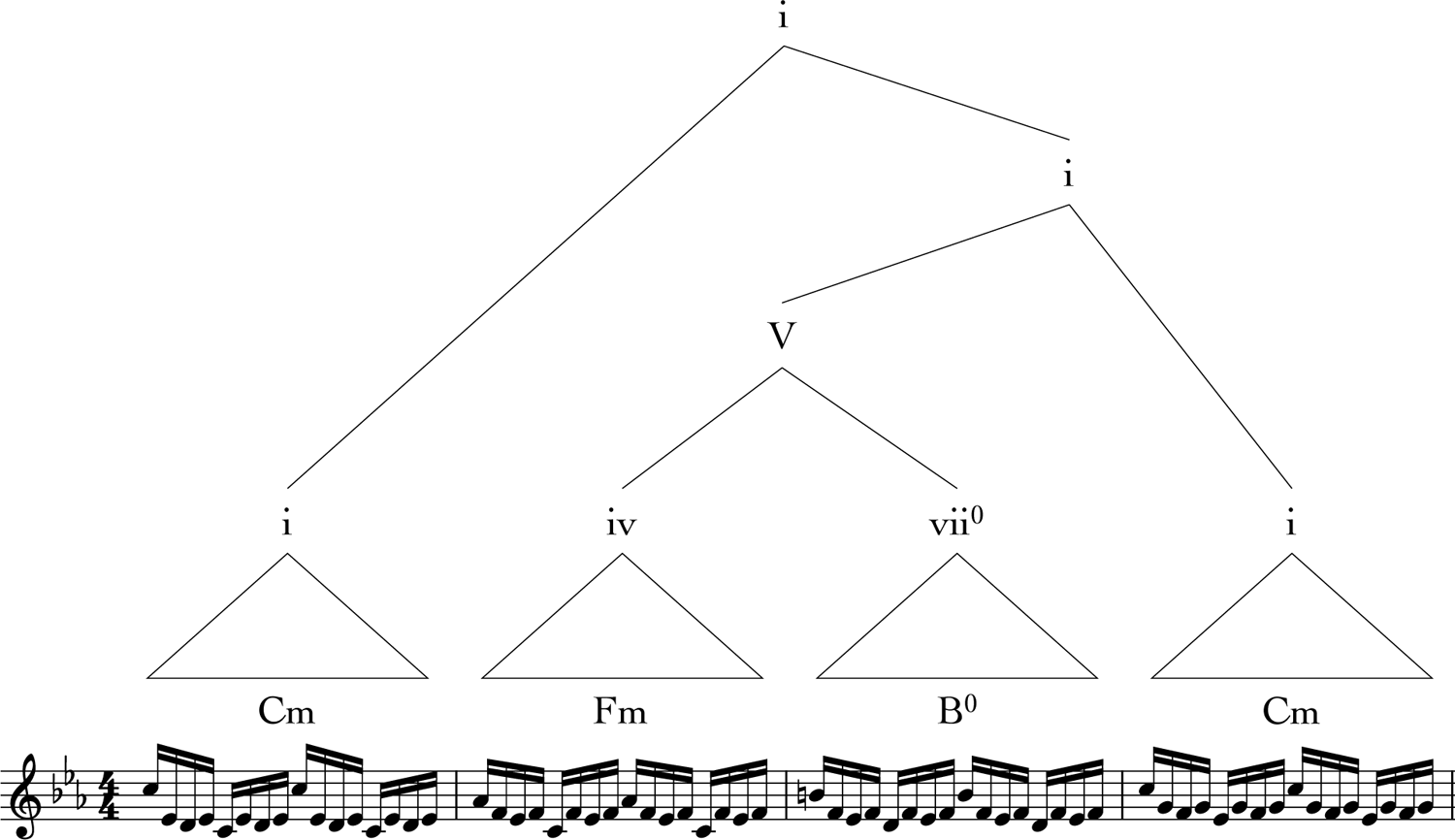
Example tree annotation of a musical phrase reveals hierarchical syntactic structures. The phrase shows the right-hand part of the opening of Bach’s Prelude in Cm, BWV 87. The leaves of the tree (i.e., the elements at the lowest level) correspond to the *terminals*, which constitute in this case the sequence of chords that characterize this musical segment, in analogy to the word sequence in a sentence.

Hierarchical analyses are commonly used in music analysis (Lerdahl & Jackendoff, 1983b; Narmour, 1983). The concrete instances of the grammar are style dependent (Katz, 2017; Rohrmeier, 2011, 2020a; Rohrmeier & Moss, 2021; Steedman, 1984). Thus, exploring neural responses to such music-theoretical structures require the formalism for the specific musical idiom. For this purpose, we adopt a syntactic formalism describing tonal harmony in common-practice Western classical music (Rohrmeier & Neuwirth, 2015).

In the following, we will define the quantification of embedding depths based on a syntactic formalism provided an expert analysis of a musical piece, so that these estimates can then be tested as predictors for brain responses. The formalism operates at the level of symbolic chord representations, that is, it represents the musical surface as a sequence of chord symbols. A rule is a function that transforms a chord symbol into a pair of other chord symbols, analogously to how a linguistic generative rule expands, for example, a (transitive) verb-phrase into a verb plus a noun-phrase (VP → V + NP) (Chomsky, 1957). The formalism allows for two distinct kinds of generative rule: *prolongations* and *preparations.* These rule types are detailed in the following and exemplified in Figure 3. In the formalism, each kind of rule can be applied to a plethora of musical chords and sequences.

**Figure 3.**
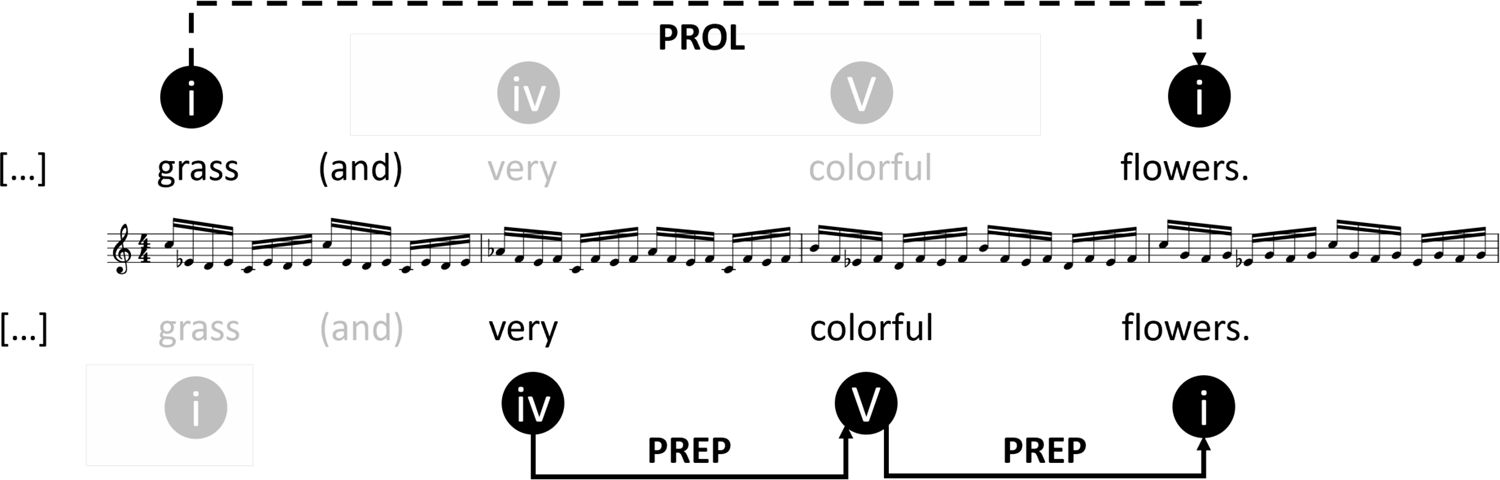
Examples of prolongations and preparation rule applications in music. The musical phrase is the same as shown in Figure 2. For illustration purposes, a linguistic example (“[I looked out of the window and saw] grass and very colourful flowers*”*) is also provided that roughly matches the dependency structure of the musical phrase.

*Prolongations* extend an event through the recursive generation of a duplication of a given chord: *A → A A*, where both instances serve the same harmonic function. To some extent, prolongations can be understood as a musical analogue to coordination in language. Both a musical as well as a linguistic example can be found in Figure 3. Note that whilst prolongation extends an event by duplication, the resulting pair of events may not have to be exactly identical on the surface. This may be because the same chord may have a different surface spelling or because both language as well as music features the possibility of substitutions. For example, in Figure 3, *‘grass’* and *‘flower’* are two different words fulfilling the same direct-object function, resulting in a ‘temporally prolonged’ object. Similarly, musical styles characterize specific kinds of substitutions (Cecchetti, Herff, Finkensiep, Harasim, & Rohrmeier, 2022; Rohrmeier, 2020a; White & Quinn, 2018).

The top half of Figure 3 highlights a case of prolongation. Through the application of a prolongation rule two tonic regions are generated that stand in a prolongation relationship to each other (also see Figure 2, the root tonic (i) is split into two tonic regions (i)). In Figure 3, this prolongation relationship between the two tonic (i) regions is indicated by the dashed line connecting Bars 1 and 4 of the musical example, which instantiate the same underlying harmony in two different ways. A key cognitive demand of prolongation is predicted to be the memory retention of the initially prolonged harmony until it, or a substitution of it, are encountered again (see Gibson, 1998, 2000).

*Preparations* elaborate an event *‘A’* by generating an event ‘*D’* that establishes an expectancy towards *‘A’: A → D A. Preparations* can be understood as a musical analogue to a linguistic rule such that the left child implies a specific word class to come as the right child, and the sentence can only be considered grammatical once a word fulfilling the implication is encountered. In the musical case, preparations are best exemplified by the relationship between a dominant and its tonic, or between a subdominant and a dominant – the first harmony in each pair generating strong expectancies towards the second. Looking at Figure 3 from a generative perspective, the 3rd bar (‘V’ harmony) of was generated through the application of a preparation rule to the final tonic (‘i’) of the phrase, and the 2nd bar of the example (‘iv’ harmony) was generated by applying a preparation rule to the newly generated ‘V’. This generation process can be seen in the tree annotation in Figure 2, by following the branches from the top to the bottom. In Figure 3, the preparation dependency relationships are indicated with solid arrows at the bottom of the figure, whereby the subdominant ‘iv’ “prepares” the following dominant ‘V’, which in turn “prepares” the following tonic ‘i’. A key cognitive demand of preparation is predicted to occur during structural integration when the dependency is resolved (see Gibson, 1998, 2000).

Both kinds of rules, prolongation, and preparation, can be recursively applied to generate a musical phrase with a complex musical structure. Importantly, since musical performance is a temporal activity, the presentation of music is inherently incremental. When listeners are presented with music, from beginning to end, sometimes they will be at a point in the musical piece where a prolongation or preparation dependency is open and not yet fulfilled. Note that this is fundamentally the same in hierarchical syntactic models of language. For example, if the example sentence in Figure 3 would be interrupted after *‘very’* (i.e., *‘I looked out of the window and saw grass and very…*’) readers or listeners would likely identify the sentence to be lacking a grammatically acceptable ending. Analogously, if the music would stop after bar 3 (‘V’) listeners would also tend to experience the lack of a concluding tonic (Herff, Harasim, et al., 2021; Pearce, 2018; Sears, Spitzer, Caplin, & McAdams, 2020). Consequently, at each given point in time, the number of open prolongation and preparation dependencies may vary. Figure 4 shows this for the same excerpt introduced in Figures 2 and 3. This example is fairly short and simple; however, over the course of a whole musical piece lasting minutes, the number of prolongation and preparation dependencies that are open at any given time can fluctuate widely. Whenever a harmony is presented that suggests the opening of a new relationship, the count for this particular type of dependency increases and whenever a harmony is presented that closes a pre-existing open dependency, the count for this type of open dependencies decreases.

**Figure 4.**
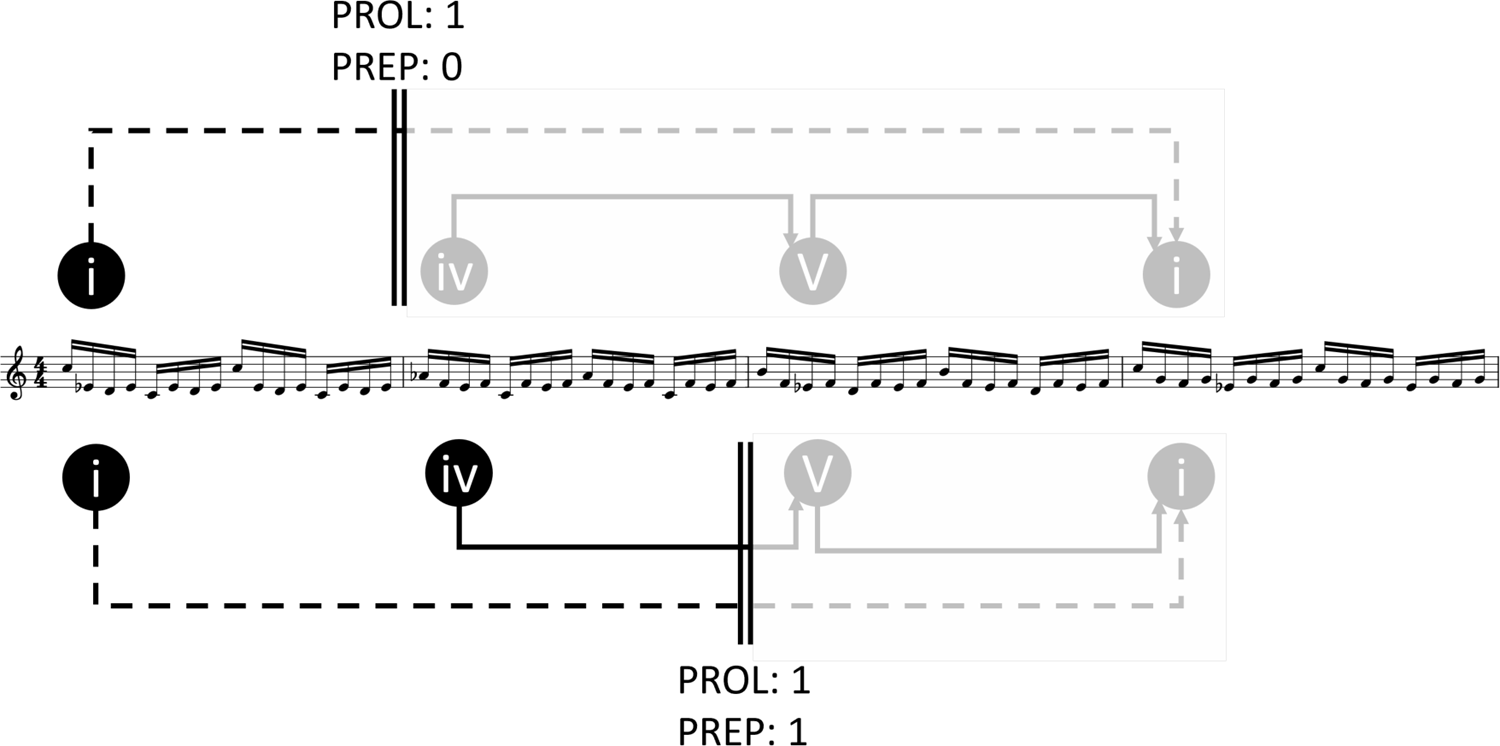
Illustrates the number of open prolongation and preparation dependencies encountered at two different time points (indicated by the double vertical lines above and below the score) whilst listening to the music. Based on the music-theoretical framework adopted here, listeners would find the music in a state of one open prolongation and no open preparation by the end of the first bar (top). Around the end of the second bar (bottom), listeners would perceive the piece in the state of one open prolongation as well as one open preparation dependency.

Processing models based on hierarchical syntax, within and outside of the field of music, predict that such open dependencies pose additional processing demands. This has seen widespread support, for example in the form of additional working memory demands (Badre, 2008; Friederici & Singer, 2015; Gernsbacher, 1997; Gibson, 1998; Gibson & Thomas, 1999; Jeon & Friederici, 2015; Kaan, Harris, Gibson, & Holcomb, 2000; King & Kutas, 1995; Lewis et al., 2006; Li & Zhou, 2010; Ma et al., 2018a, 2018b; Ma et al., 2022; Makuuchi, Bahlmann, Anwander, & Friederici, 2009; Makuuchi & Friederici, 2013; Meyer et al., 2015; Meyer, Obleser, & Friederici, 2013; Opitz & Friederici, 2007; Phillips, Kazanina, & Abada, 2005; Stromswold, Caplan, Alpert, & Rauch, 1996; Vos, Gunter, Kolk, & Mulder, 2001). Conveniently, the formalism summarised above allows us to precisely quantify the number of open dependencies at every moment in time within a musical piece (see Figure 4). Consequently, in the present study, we aim to utilise the fluctuation of open prolongation and preparation dependencies throughout a piece as a continuous predictor of theta activity, in an attempt to both test the predictions of this particular formalism and identify brain regions involved in the processing of the hierarchical dependencies it predicts.

### Aim and Hypotheses

Here, we aim to test whether a music-theoretical analysis of a musical piece, formalised as a hierarchical syntactic structure, carries predictive value for listeners’ theta activity when listening to the same piece. This allows us to critically test the predictions of the music-theoretical model, whilst identifying potential brain regions reflecting syntactic processing in music. The aforementioned studies showing that theta power is stronger in pronouns with embedded compared to non-embedded antecedents (Meyer et al., 2015) (also see Figure 1), that theta can be associated with higher memory load (Jensen & Tesche, 2002), combined with the assumption that an increase in open dependencies during the parsing of syntactic structures should be associated with rising demands on the memory system (Lewis et al., 2006), allows us to generate direct predictions for the musical case. Specifically, work in psycholinguistics suggests that (at least) two types of cognitive operations are required to implement syntactic processing (Gibson, 1998, 2000): (1) memory storage for retaining representation of past or future events in memory, and (2) integrating the currently-parsed event into an existing partial parse (structural integration). Extending this framework to the musical domain, we suggest that (1) prolongations are predominantly associated with memory storage demands extending from the time the prolonged harmony is first encountered to the time the prolongation is completed by a return to the same harmony or a substitute thereof leading to the prediction of increases theta power with increasing number of open prolongation dependencies. Preparations, on the other hand, is predominantly associated with the structural integration of the events that resolve the dependency, so that greater activation is predicted during integration, that is increased theta power as the number of open preparations decreases.

As some prior literature has shown auditory envelope tracking in the theta band, we also assess whether evidence for the observed effects remain after controlling for envelope tracking within the theta frequency band. In a final step, we assess whether musical expertise mediates any of the effects observed.

## Method

Here, we used the same dataset employed in a previously published MEG study (Bonetti, Brattico, et al., 2021). In the current work, the pre-processing pipeline for the MEG data and the source reconstruction algorithms are similar to the ones described in Bonetti et al. (2021), however, there are a few differences that are described in the following sections. Moreover, additional steps were taken to obtain the music-theoretical annotation of the musical piece, extract theta band envelope, and construct the statistical models attempting to link music theoretical predictions with brain responses. The MEG data is available upon reasonable request, whereas all analyses as well as the full tree annotation of the stimulus pertinent to the present study can be found on https://osf.io/h8x47/.

### Participants

The dataset consists of 65 right-handed and 3 left-handed participants, demographics of the sample are summarized in Table 1. Participants were recruited from three different musical expertise levels. This allows to control for potential expertise effects, as prior studies have demonstrated that brain responses to musical structures can be modulated by levels of musical expertise (Koelsch, Schmidt, & Kansok, 2002). All participants were required to be fluent in English and have normal or corrected-to-normal hearing and sight. Musicians were mainly recruited among bachelor’s and master’s students at the Royal Academy of Music Aarhus/Aalborg (RAMA). Additional musicians were recruited by posting messages on social networks. Non-Musicians were from the bachelor’s and master’s students at Aarhus University and via posting messages on social networks. Following this procedure allowed us to recruit participants with approximately the same age and socio-economic and educational background.

**Table 1.** Participants’ demographic information

| **Group** | ***N*** | **Self-reported**<br>***M<sub>Age</sub> (SD<sub>Age</sub>, Range<sub>Age</sub>)*** | **Self-reported**<br>****Gender**** |
| --- | --- | --- | --- |
| **Non-Musicians** | 24 | 25.46 (3.48, 21-35) yrs. | 12 female, 12 male |
| **Musicians** |  |  |  |
| Classical Pianists | 23 | 24.83 (4.10, 18-34) yrs. | 10 female, 13 male |
| Non-Pianist | 21 | 24.29 (5.02, 19-42) yrs. | 11 female, 10 male |
| **Total** | 68 | 24.88 (4.17, 18-42) yrs. | 33 female, 35 male |

### Stimuli

The stimulus was an adaptation of the Prelude in C minor BWV 847 composed by Johann Sebastian Bach, a piece firmly rooted within the Western tonal-music tradition, and as a consequence, within the scope of the formalization of musical syntax (Rohrmeier, 2011; Rohrmeier & Neuwirth, 2015). The piece has a specific compositional and textural setup such that for most of its duration both hands play the same pattern constituting a verticalization of the underlying harmony. Therefore, for the purposes of the original study (Bonetti, Brattico, et al., 2021), it was possible to transcribe the piece into a reduced monophonic version, in which only musical notes notated in the score for the right-hand piano part were presented, and occasional gaps in the right-hand part were filled with left-hand material with the necessary register or voicing adaptations. Crucially for our analysis, the harmonic implications that are the foundation of the syntactic dependencies of the piece remained intact. Additionally, the original study required each note to have approximately, but not exactly, the same length. Consequently, each sounded sixteenth note was quantized to last 250ms on average, plus a random deviation of up to ± 30ms.

While most of the piece is characterized by a uniform texture of repeated sixteenth-notes, bar 34 exhibits a sparser texture with longer sounded notes that were modified in order to preserve the sixteenth-note pattern. A MIDI version of the final stimulus was synthesised in the default piano timbre of Finale (MakeMusic, Boulder, CO). The volume was adjusted individually to be 50 dB above each participants’ minimum hearing threshold. The score, midi files, as well as synthetised wave file of the stimulus can be found in https://osf.io/h8x47/.

The stimulus, as presented to the participants, was analysed and annotated by the last author using the aforementioned formalism through a dedicated web-app (Harasim et al., 2020). For this, chord labels were attached to the chords formed or implied in the monophonic notated score. Within the formalism, these chord labels specify functional harmonies. Syntactic relations between chord labels were then encoded as a derivation tree under the grammar proposed by Rohrmeier & Neuwirth (2015). At the location of each harmonic chord label (the terminals of the derivation), the number of open *Prolongations* and *Preparations* was measured as previously described. Specifically, for every non-terminal node of the analysis, the count of open dependencies of the type corresponding to the rule that is applied to that non-terminal is increased by 1 in the timespan between the terminals that label the left and right child of the node, respectively. Furthermore, each prolonged instance of the left child of a preparation also contributes to increase by 1 the count of open preparations until the target (right child) of the preparation is encountered. Note that musical annotations are more ambiguous than linguistic annotations, with some freedom of the annotators to interpret specific sections. To have a rough measure of inter-annotator agreement for harmonic analyses, the third author independently performed the same annotation task. The number of open *Prolongations* and *Preparations* derived from the two annotations were Spearman correlated with *r* = .73 and *r* = .93, respectively. The full final tree annotation can be found in https://osf.io/h8x47/.

Note that, since the stimulus presented to participants only contained a monophonic stream, harmonies were not directly present in the form of chords, but rather implied by means of latent polyphony (Aldwell et al., 2018; Davis, 2006; Finkensiep & Rohrmeier, 2021; Holleran, Jones, & Butler, 1995). Furthermore, since the stimulus only comprised the right-hand part in a piano texture, some harmonic progressions, that would normally be strongly instantiated or reinforced by the bass in the left-hand, were less clearly presented to the participants. This makes the present a study a conservative attempt to link hierarchical music theoretical annotations with brain responses, as in most cases of idiomatic common-practice music the harmonic texture is fully specified or at least less underspecified than it is in the present stimulus.

### Neural data acquisition

The MEG data was acquired utilizing an Elekta Neuromag TRIUX MEG scanner with 306 channels (Elekta Neuromag, Helsinki, Finland) in a magnetically shielded room at Aarhus University Hospital (AUH). The data used in the study was collected at a sampling rate of 1000 Hz with an analogue filter of 0.1 to 330 Hz. The participants’ head shapes and the positions of four Head Position Indicator (HPI) coils were recorded using a 3D digitizer (Polhemus Fastrak, Colchester, VT, USA) in relation to three anatomical landmarks (left and right preauricular points, and nasion). This recording was later utilized to co-register the MEG data with the MRI scans. During the MEG recordings, the HPI coils continuously monitored the head’s location and this information was used to correct for participants’ movement in the scanner. Additionally, two sets of bipolar electrodes were used to record the participants’ cardiac rhythm and eye movements. This enabled the removal of electrocardiography (ECG) and electrooculography (EOG) artifacts during the pre-processing of the data.

The MRI scans were conducted on a CE-approved 3T Siemens MRI scanner at AUH and included structural T1 images with a resolution of 1.0 x 1.0 x 1.0 mm and the following sequence parameters: echo time (TE) = 2.61 ms, repetition time (TR) = 2300 ms, echo spacing = 7.6 ms, reconstructed matrix size = 256 x 256, bandwidth = 290 Hz/Px. The MEG and MRI recordings were conducted on separate days.

### MEG data pre-processing

The raw MEG sensor data, consisting of 204 planar gradiometers and 102 magnetometers, was initially pre-processed using MaxFilter (Taulu & Simola, 2006) to reduce external interference. We applied signal space separation using the following MaxFilter parameters: spatiotemporal signal space separation (SSS), down-sampling from 1000Hz to 250Hz, and movement compensation using cHPI coils (with a default step size of 10 ms). The correlation limit between inner and outer subspaces used to reject overlapping intersecting inner/outer signals during spatiotemporal SSS was set to 0.98. The data was subsequently converted into Statistical Parametric Mapping (SPM) (Henson & Friston, 2007) format and further pre-processed and analysed using a combination of in-house-developed codes (LBPD, available at https://github.com/leonardob92/LBPD-1.0.git) and the Oxford Centre for Human Brain Activity (OHBA) Software Library (OSL) (https://ohba-analysis.github.io/osl-docs/) (Woolrich, Hunt, Groves, & Barnes, 2011), which is a free software that builds on the Fieldtrip (Oostenveld, Fries, Maris, & Schoffelen, 2011), FSL (Woolrich et al., 2009) and SPM toolboxes. All of this was done using MATLAB (MathWorks, Natick, MA, USA). The MEG data was carefully examined for large artifacts, and any identified artifacts were removed using the OSLview tool. The amount of data removed in this process was less than 0.1% of the total collected data. To remove the interference of eyeblinks and heart-beat artifacts from the brain data, we employed independent component analysis (ICA) (Mantini et al., 2011). The process involved decomposing the original signal into independent components, identifying, and removing the components that captured eyeblink and heart-beat activities, and then reconstructing the signal using the remaining components.

### MEG source reconstruction

The Polhemus head shaped data, combined with the three fiducial points acquired during the MEG recording was used to co-register each individual T1-weighted MRI scan to the standard Montreal Neurological Institute (MNI) template brain through an affine transformation.

The co-registered images where then used to reconstruct the sources of the signal recorded with the MEG scanner. To this end, we used a beamforming algorithm, independently for each participant (Hillebrand & Barnes, 2005; Huang, Mosher, & Leahy, 1999; Brookes et al., 2016).

This technique consists of two main steps: (i) designing a forward model and (ii) computing the inverse solution.

The forward model is a theoretical model that represents each brain source as an active dipole (brain voxel) and describes how the unitary strength of each dipole would be reflected by the MEG sensors. In this study, we used magnetometer and gradiometer channels and an 8-mm grid, which resulted in 3559 dipole locations (voxels) within the whole brain. We calculated the forward model using the widely used “Single Shell” method (Nolte, 2003). In the three cases where structural T1 data was not available, we performed the forward model calculation using a template (MNI152-T1 with 8-mm spatial resolution).

We then calculated the inverse solution using beamforming. This involves applying a set of weights sequentially to the source locations in order to isolate the contribution of each source to the activity recorded by the MEG channels, at each timepoint of the recorded brain data. The 3559 brain sources were then projected on the widely used Automated Anatomical Labelling (AAL) atlas (Tzourio-Mazoyer et al., 2002) by utilizing principal component analysis (PCA). The AAL atlas contains 116 structure descriptions; however, here we focused only on the 90 brain regions, excluding the cerebellum.

For further information regarding pre-processing of the MEG data and source reconstruction, please refer to the following works (Bonetti, Brattico, et al., 2022; Bonetti, Bruzzone, et al., 2021; Bonetti, Carlomagno, et al., 2022; Fernández-Rubio, Brattico, et al., 2022; Fernández-Rubio, Carlomagno, Vuust, Kringelbach, & Bonetti, 2022).

### Procedure

After providing informed consent, participants were comfortably seated in the MEG scanner. First, a 10-minute-long resting state session was recorded, in which participants were instructed to relax, but not to close their eyes. The resting state session served to familiarise participants with the environment of an MEG scanner, and was used as a baseline condition for a function connectivity analysis reported elsewhere (Bonetti, Brattico, et al., 2021). Afterwards, participants were presented with the musical stimulus and instructed to attentively listen to the music, trying to memorize its structure and sounds as much as they could. Each participant was presented four times with the musical stimulus, with self-paced pauses between each presentation. Afterwards, participants were presented with a recognition task that required participants to distinguish the previously presented musical stimulus from previously non-presented foils in the same musical idiom (see Bonetti et al., 2020, for further detail). The task served as a test of whether participants attentively listened to the musical piece. All experimental procedures complied with the Declaration of Helsinki – Ethical Principles for Medical Research and were approved by the Ethics Committee of the Central Denmark Region (De Videnskabsetiske Komitéer for Region Midtjylland, Ref 1-10-72-411-17).

### Statistical Analysis

#### Preprocessing

To obtain theta activity, we extracted power in the 3-7Hz band from the broadband MEG data reconstructed in source space using the ‘eegkit’ package (Helwig, 2018) implemented in R (RCoreTeam, 2021). We then extracted the Hilbert amplitude envelope of the theta band power using the ‘seewave’ package (Sueur, Aubin, & Simonis, 2008). To assess whether the music-theoretical concept of prolongation and preparation carries predictive value for brain activity, we averaged the envelope of the theta band activity across the corresponding times points of the four repetitions of the piece, but separately for the 90 brain regions and each participant. Whenever the harmonic chord label in the piece changed (based on the harmonic annotation of the piece provided by two expert annotators) a new stimulus locked marker was set, resulting in a total of 67 marker positions. Theta envelope in a 1000ms long segment starting from marker onset was then averaged separately for each participant and their respective brain regions. The original data set did not record data for the final chord, as it was irrelevant to the original research question (see Bonetti et al., 2020 for further detail). Consequently, a total of 403920 segments were analysed, comprising 66 harmonic marker positions, 68 participants, and 90 brain regions. Additionally, the piece exhibits a shift in harmonic rhythm starting in bar 28. Specifically, the rate at which the harmonic chord labels change is substantially faster towards the end of the piece (3.8 per bar on average), compared to the beginning (1 label per bar). This is important to note, as harmonic density might influence listeners’ ability to extract underlying harmonies and relationships between harmonies in this monophonic rendition of the piece. To account for this, we conducted two separate analyses on slow (bars 1 to 27) and fast harmonic rhythm (bars 28 to 38) sections in the piece (preferably, we would have included harmonic rhythm as an interactive term in the model rather than splitting the data; however, we were simply lacking the computational power to do so). We report the results separately for the slow- and fast-harmonic-rhythm sections for interested readers; however, for the purposes of the present work, we interpret the results jointly for the entire piece as we are not concerned with variable difficulty.

#### Identifying Brain Regions of Interest

We implemented Bayesian Mixed Effects Models that used the number of open prolongations and preparations identified by the expert annotators to predict average theta envelope in each segment. The models were implemented in R (RCoreTeam, 2021) using the brms package (Bürkner, 2017). All continuous variables were standardised to have a mean of 0 and a standard deviation of 1, and the model was provided with a random effect for participant. Each model ran on 4 chains, was initialised with 0s, and was provided with a weakly informative prior in form of a t-distribution with mean 0, a standard deviation of 1, and 3 degrees of freedom (Gelman, Jakulin, Pittau, & Su, 2008), a prior commonly used in recent music perception studies (Beveridge, Cano, & Herff, 2021; Herff, Cecchetti, Taruffi, & Déguernel, 2021; MacRitchie, Breaden, Milne, & McIntyre, 2020; Milne, Dean, & Bulger, 2021; Smit, Dobrowohl, Schaal, Milne, & Herff, 2020). We then conducted hypothesis tests on the posterior of the fitted models for effects of prolongation and preparation in each region. We report the model coefficients (*β*) relevant to the specific regions, the estimated error of this coefficients, as well as the evidence ratio in favour of a given hypothesis. We consider regions of interest only those brain regions that show prolongation or preparation effects with evidence ratios of at least 19 (Milne & Herff, 2020). Due to the size of the data frame and the computational demand of the models, this analysis was conducted separately for the left and right hemisphere. After brain regions of interest were identified, we further examined the effects to ensure their reliability.

#### Consolidating The Results

To critically test whether the observed predictive effects of the theoretical concepts of prolongations and preparations was an artifact of purely acoustic features of the music, we conducted two additional tests. First, we implemented an additional Bayesian mixed effects model that only included data from the previously identified regions of interest. In this model, the theta envelope of the audio — segmented and extracted in same fashion as the theta envelope from brain data — was included as an additional predictor. The presence of this predictor and its interaction with each brain region in the model, as well as its low correlation with the prolongation and preparation predictors (*r* = .161 and *r* = −.052 respectively) ensure that any remaining predictive value of prolongation and preparation cannot be explained by audio envelope tracking instead. This is an important consideration, as audio-envelope tracking is commonly observed in neural activity in the auditory domain (Aiken & Picton, 2008; Pérez et al., 2022) and in music in particular (Di Liberto, Pelofi, Shamma, & de Cheveigné, 2020; Herff et al., 2020; Miyazaki et al., 2013; Zuk et al., 2020). Furthermore, to ensure that the predictive value of prolongation and preparation resides in the precise link of harmonic chord labels – and the prolongation and preparation dependencies they imply – to a specific temporal point, rather than the general distribution of labels, we randomly shuffled the prolongation and preparation predictors 10000 times. After each shuffle, which disrupted the link between predictor value and music, we ran the model again and recorded the observed effect sizes. Only effects with an evidence ratio of at least 19 whilst controlling for audio envelope tracking and whose effect sizes exceeded the magnitude of 95% of this random effect distribution were considered reliable effects and are reported in the results.

#### Musical Expertise

In a last step, we assessed whether musical expertise mediates the predicative value of music theoretical prolongation and preparation for theta envelope in any of the brain regions identified in the previous step. For this, we implemented another Bayesian Mixed Effects model, identical to the ones describe above, but including an additional region of interest*musical expertise interaction term, with musical expertise being modelled as a categorical factor with levels Non-Musician, Pianist, and Non-Pianist Musician. We then report in which regions predictive values of prolongation and preparation differed reliably in Pianist or Non-Pianist Musician from Non-Musicians.

## Results

All fitted models can be found in the online supplement at https://osf.io/h8x47/. Table 1 reports those brain regions for which prolongation or preparation carry predictive value for theta fluctuation in the part of the piece with the slow harmonic rhythm. Table 2 reports the same for the section of the piece with the fast harmonic rhythm.

**Table 2.**
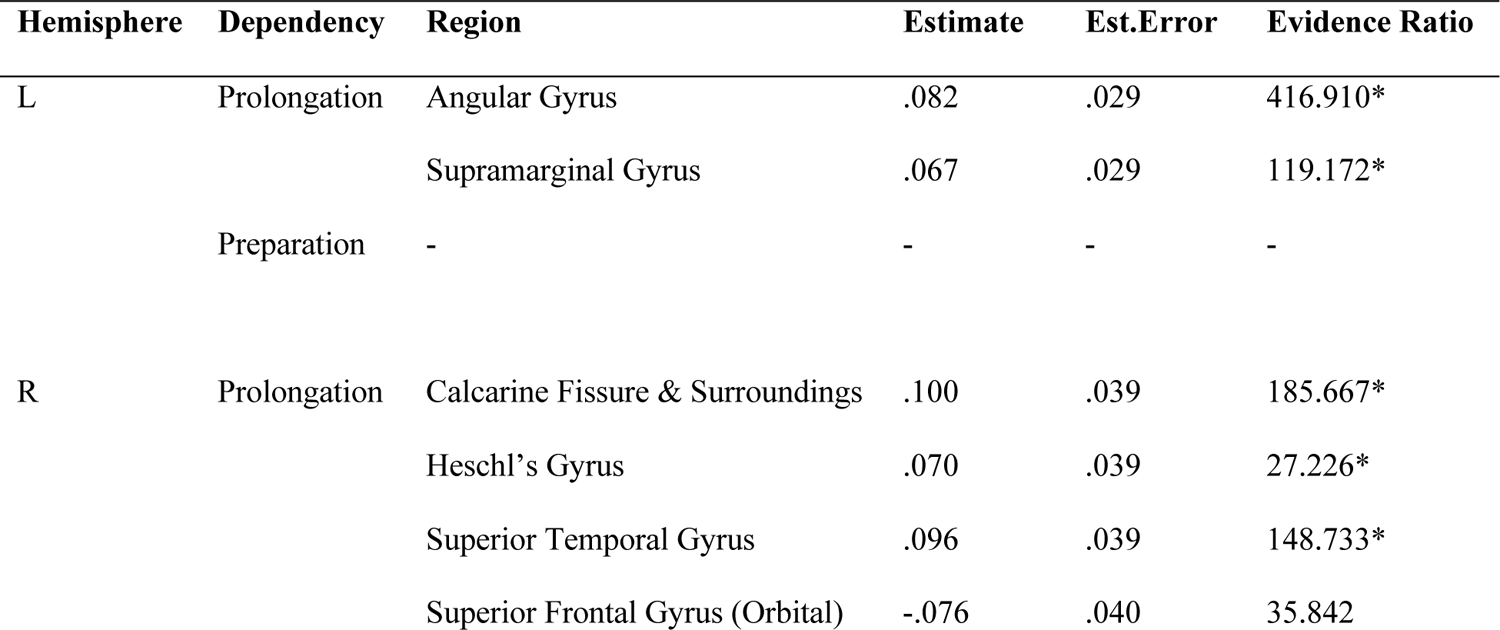
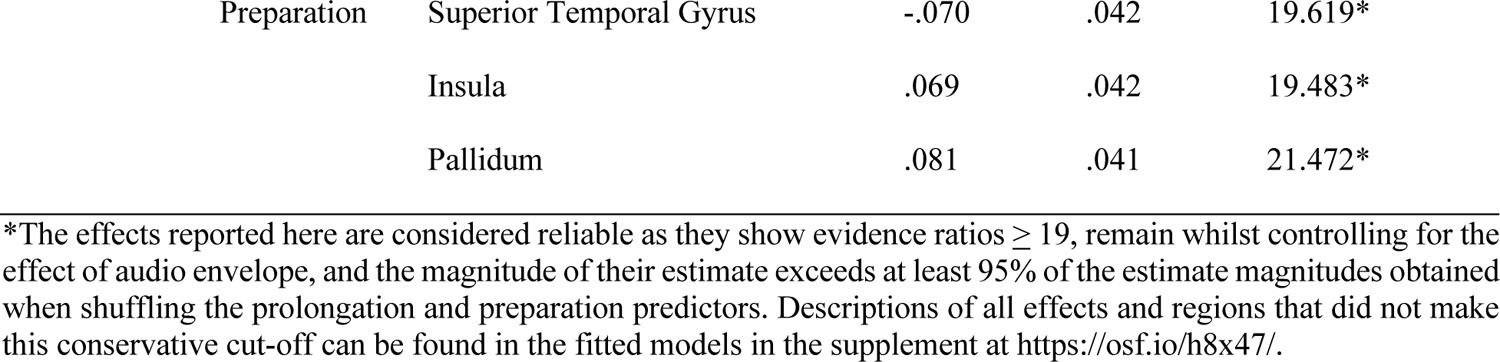
Reliable effects in the slow harmonic rhythm section.

**Table 3.**
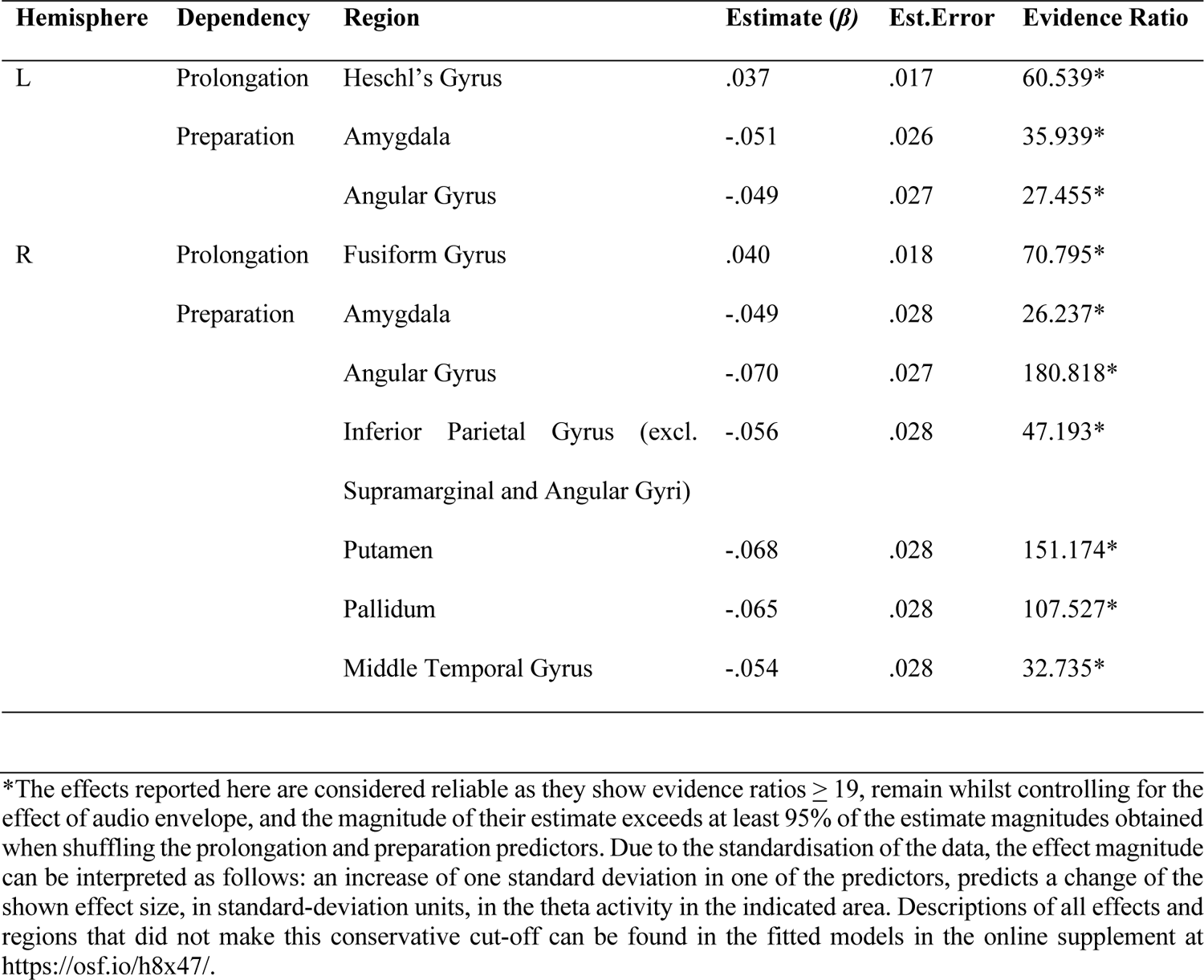
Reliable effects in the fast harmonic rhythm section.

Figure 5 shows cortical brain areas in which the two music-theoretically-informed predictors carry predicative value for brain theta envelope across the musical segments with fast and slow harmonic rhythm, displayed for preparation and prolongation separately. As mentioned above, only those effects are shown that are classified as reliable, that is those that have at least an evidence ratio of 19 whilst controlling for the effect of audio envelope and exceed 95% of the chance distribution. We plot the results in the AAL atlas that we also used for localisation. Whilst we plot the results in a common space, the analysis was conducted in the original recorded space of each participant. A breakdown for the results for every region can be found in supplement (https://osf.io/h8x47/). The figure highlights the involvement of Inferior Parietal, in particular for preparations, as well as additional areas in Temporal and Frontal areas for prolongation. The figure also shows a tendential right lateralisation.

**Figure 5.**
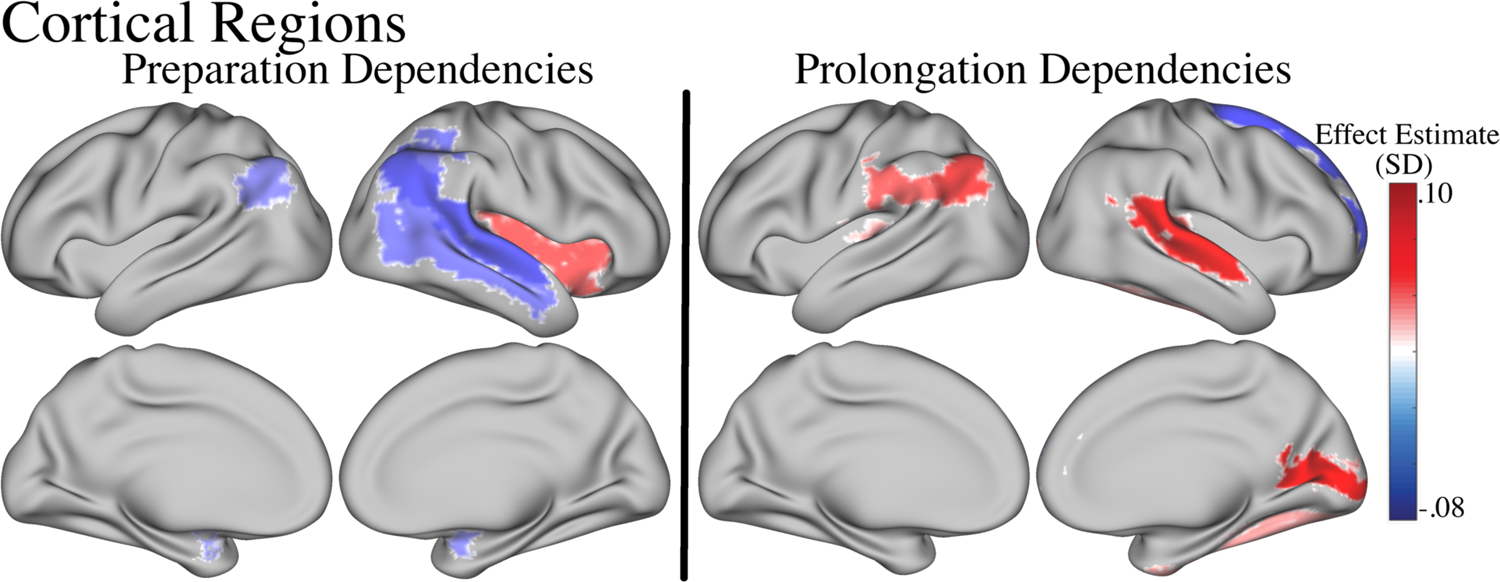
Overview of cortical brain areas in which the music theoretically informed preparation (left) and prolongation (right) predictors carry reliable predictive value for theta brain activity.

Figure 6 shows subcortical areas in which preparation and prolongation carry predictive value for Theta envelope, plotted on the Freesurfer automatic subcortical segmentation of a brain volume (Fischl et al., 2002). We did not observe strong evidence for prolongation dependencies to carry reliable predictive power for theta envelope, however, preparation carried predictive value for theta envelope fluctuation in multiple subcortical areas: bi-lateral Amygdala and the right Putamen as well as Pallidum.

**Figure 6.**
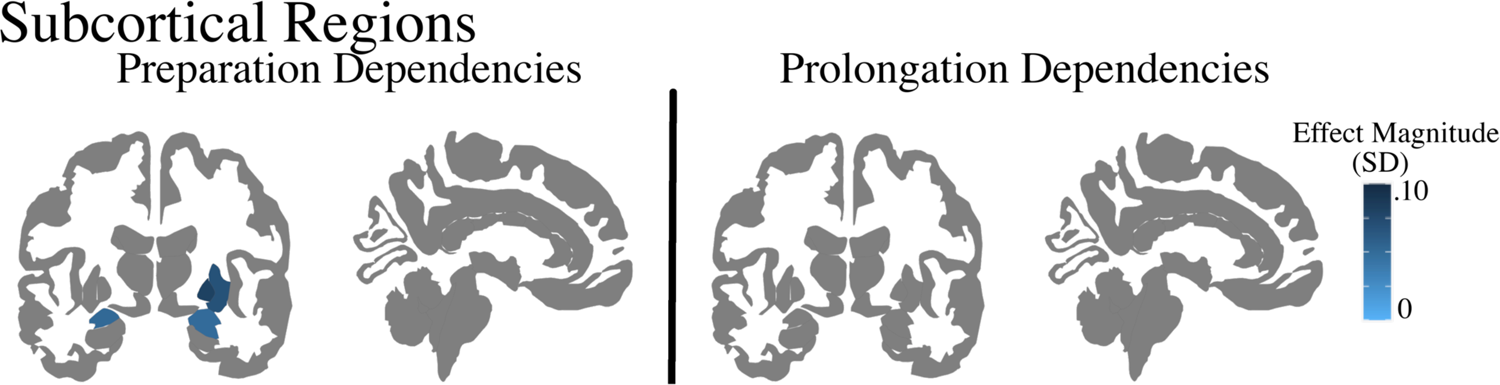
Overview of subcortical brain areas in which the music theoretically informed preparation and prolongation predictors carry reliable predictive value for theta brain activity. We plot effect magnitude due to the conflicting directionality in the Putamen between in the slow and fast harmonic rhythm. Preparation, but not prolongation, carries predictive value for theta brain activity in subcortical regions.

### Musical Expertise

As shown in Figure 7 top-row, the left Superior Temporal Gyrus was the only area in which we obtained some evidence that the predictive value of music-theoretical preparation was mediated by musical expertise (*Estimate* = −.16, *Est.Error* = .09, *Evidence Ratio* = 21.56*), indicating larger effect magnitudes of preparation on theta band activity in non-musicians compared to musicians.

**Figure 7.**
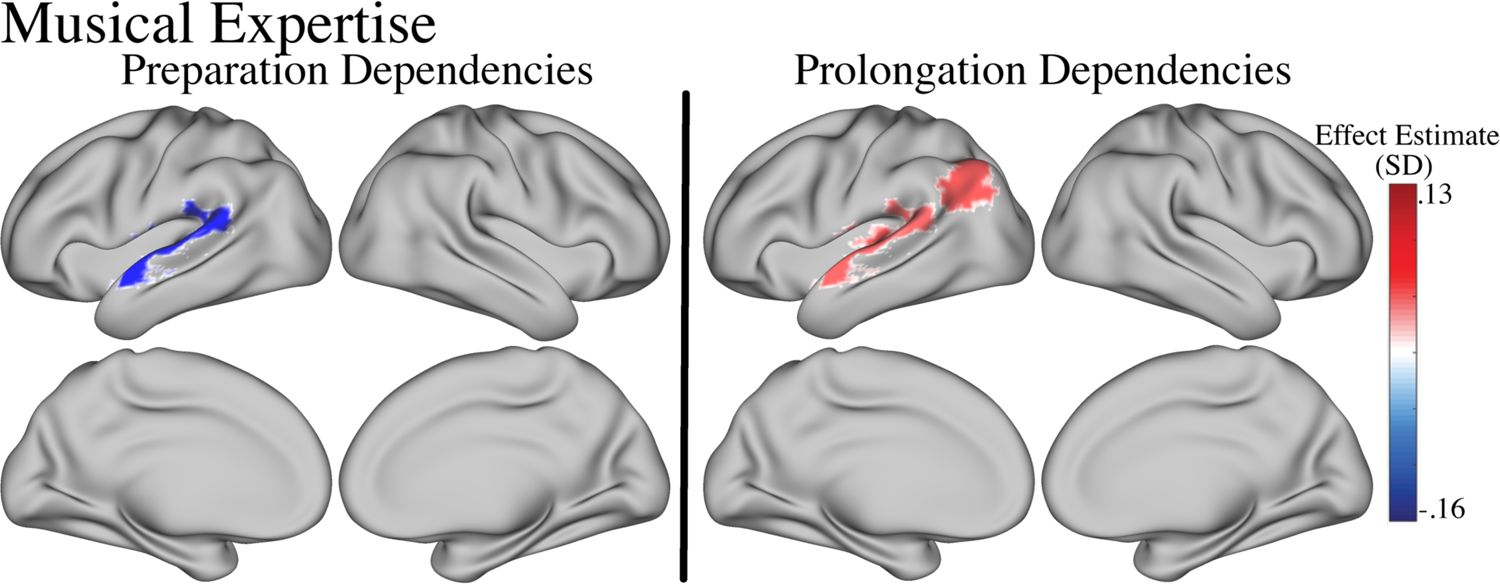
Overview of brain areas in which musical expertise affects the predictive value that music theoretically informed preparation and prolongation predictors carry for theta brain activity.

We also observed strong evidence that musical expertise mediated the predictive value of music-theoretical prolongation dependencies in the left Heschl’s Gyrus (*Estimate* =.13, *Est.Error* = .09, *Evidence Ratio* = 75.71*), left Superior Temporal Gyrus (*Estimate* =.11, *Est.Error* = .06, *Evidence Ratio* = 29.27*), as well as the left Angular Gyrus (*Estimate* =.12, *Est.Error* = .07, *Evidence Ratio* = 22.35*), with larger effects of prolongation on theta band activity in non-musicians compared to musicians in all regions. This is visualised in Figure 7, bottom-row.

## Discussion

Hierarchical syntactic models of language predict increasing processing demands with deeper embedding depths, which can be observed in the theta band (Meyer et al., 2015). Here, we tested whether a similar phenomenon can be observed with musical stimuli. We used an expert music-theoretical annotation of a musical piece based on a hierarchical syntactic model of harmonic relationships (Rohrmeier & Neuwirth, 2015) to continuously quantify embedding depths of two types of syntactic embeddings: prolongations and preparations. Using fluctuations in these two types of embeddings to predict brain theta activity, we showed that embedding depth as formalised by a hierarchical syntactic model of harmony carries reliable predictive value for brain responses, and that this effect is affected by musicianship. In the following, we discuss the findings for prolongation and preparation and propose explanations at the computational and algorithmic level (Marr, 1982) for the observed brain activity.

*Prolongation* can be loosely understood as a musical analogue to the linguistic coordination (Granroth-Wilding & Steedman, 2014; Steedman, 1984) investigated in Meyer et al. (2015). Meyer et al. observed an increase in theta activity when the antecedent of a pronoun was grammatically embedded compared to when it was not. Prior studies already demonstrated that theta activity can be used to mark segment boundaries of musical sections (Quon et al., 2021). We here extend this finding, by showing that continuous embedding depth in the form of open prolongation dependencies carried predictive value for theta power in the left and right Heschl’s Gryus, the left Angular Gyrus and Supramarginal Gyrus, the right Superior Temporal Gyrus and Calcarine Fissure and Orbital Part of the Superior Frontal Gyrus as well as Fusiform Gyrus. The Heschl’s Gyrus has long been associated with primary auditory processing (Da Costa et al., 2011; Warrier et al., 2009; see Saenz & Langers, 2014, for a review), for example in the form of envelope tracking (Kubanek, Brunner, Gunduz, Poeppel, & Schalk, 2013; Nourski et al., 2009). However, since we specifically controlled for envelope tracking, our results suggest that the Heschl’s Gyrus is also involved in some early structural processing. As only prolongations, and not preparations, show predictive value for theta power in the Heschl’s Gryus, a speculative explanation could be that the Heschl’s Gyrus continuously performs an identity matching for all open prolongation dependencies. Specifically, prolongation describes a dependency relationship between two events that perform the same harmonic function. This can either be a literal repetition of the initial harmony, or the instantiation of a chord that can function as a harmonic substitute. The task of testing for identical or functionally identical repetition lays well within the scope of processes previously attributed to the Heschl’s Gyrus, such as generating mismatch negativities for auditory oddballs (Opitz, Mecklinger, von Cramon, & Kruggel, 1999), tracking of musical beat (Nozaradan et al., 2017; Nozaradan, Schönwiesner, Keller, Lenc, & Lehmann, 2018), or tagging of repetitive rhythmic patterns (Herff et al., 2020; Herff, Johnson, Milne, Herff, & Krusienski, 2017). In other words, a process that can capture whether an auditory event is inconsistent with a previously established pattern (as observed in oddball paradigms), also necessarily carries information about when it is consistent (as in prolongation cases). This would also explain why preparations were not found to carry predictive value for theta-band activity in the Heschl’s Gryus, as preparations entail the expectation of a specific harmony that is different from the one inducing the expectation: identifying an occurrence of such a target harmony is a different task than that performed when faced with prolongations or oddballs. Such directed predictions as observed in the case of preparation might require additional higher order processing, beyond the primary function of the Heschl’s Gryus, such as additional memory resources or access to a lexicon.

The involvement of the Angular Gyrus in the processing of prolongation is also well within the scope of the theoretical framework that predicted higher processing demands and memory burden as embedding depth increases. The Angular Gyrus is associated with both language processing (Bemis & Pylkkänen, 2013) as well as memory retrieval (Kim, 2010; see Seghier, 2013 for review). The tight link of the Angular Gyrus with the Supramarginal Gyrus, a structure for which prolongation also carried predictive power, is associated with phonological processing (Hartwigsen et al., 2010; Penniello et al., 1995). Interestingly, we found a predominantly left-lateralised effect, similar to language processing (Oberhuber et al., 2016). This finding supports early processing similarities between language and music (Sammler et al., 2009; Y. Sun et al., 2018) which are further emphasised by the involvement of the Superior Temporal Gyrus.

The Superior Temporal Gyrus is involved in a wide variety of speech-relevant tasks and contains Wernicke’s area, a key structure to the processing of speech in particular (see DeWitt & Rauschecker, 2013, for a review). Importantly, we here only observe the number of open prolongation dependencies to predict theta activity in the right Superior Temporal Gyrus, whereas Wernicke’s area tends to be located in the left hemisphere in right-handed participants (Rasmussen & Milner, 1977). Intriguingly, the corresponding region in the non-dominant hemisphere has been previously found to be active during the processing of linguistic ambiguity (Harpaz, Levkovitz, & Lavidor, 2009; Mason & Just, 2007). The activation of the right Superior Temporal Gyrus observed in our study may then reflect ambiguity resolution in music, since – due to the lack of communicative pressure – the parsing of musical structure has been described to be even more prone to ambiguity than the parsing of linguistic sentences (Cecchetti, Herff, & Rohrmeier, 2022). This finding is also in line with much prior literature suggesting that music processing in general tends to be right-lateralised compared to language processing, which tends to be left-lateralised (Koelsch, 2009; Scharinger, Knoop, Wagner, & Menninghaus, 2022). Whether the right-lateralisation in music processing is better understood as a music specific process, or as an extension of the same network that deals with ambiguous linguistic stimuli (Faust & Mashal, 2007; Pobric, Mashal, Faust, & Lavidor, 2008) is an intriguing question for future research. As hypothesised, the effect directionality in the Heschl’s Gyrus, Angular Gyrus, Supramarginal and Superior Temporal Gyrus are all positive, indicating an increase in theta band power as the number of prolongation dependencies increases. This is in line with the general theoretical framework, suggesting that increased theta power is indicative of higher memory load, and that deeper embedding depth is indicative of higher memory load. Additionally, this is also in line with the linguistic study (Meyer et al., 2015) that served as inspiration for the present study in the musical domain.

Whilst all previously discussed areas are in line with prior literature, the effects observed in the right Superior Frontal Gyrus, Fusiform Gyrus as well as the Calcarine Fissure were unexpected and we currently do not have any compelling explanation. The larger number of participants (N = 68), and the conservative approach of combining Bayesian Mixed Effects models with additional Monte Carlo simulation, whilst controlling for the effects of envelope tracking makes false positives unlikely. It is possible that listeners engaged in visual imagination or systematic eye movement, induced by the music (Herff, Cecchetti, et al., 2021; Taruffi & Küssner, 2019). If such behaviour were to correlate with musical structure, then this might be the source of the findings in the Fusiform and Calcarine areas, which are predominantly associated with vision, including imagined visual content (Le Bihan et al., 1993). The Superior Frontal Gyrus is associated with working memory (Boisgueheneuc et al., 2006), so its involvement is not surprising. What is surprising, however, is that the effect direction is negative, indicating weaker theta band activity with increasing number of prolongation dependencies. Potentially, driven by musical structure, participants felt the urge to tap along to the music, and suppressed this impulse; this may then be related to the systematic activity observed in the orbital part of the right Superior Frontal Gyrus, which plays a large role in impulse control (Hu, Ide, Zhang, & Chiang-shan, 2016). However, both these explanations are highly speculative and would require further testing, far beyond the scope of the present study.

*Preparation,* from a music theoretical perspective, can be understood as a generative step that introduces a harmonic event with the function of eliciting a strong expectation towards a different target harmony, such as a dominant awaiting resolution to its tonic chord. Similarly to prolongations, we observe a number of brain regions in which the number of open preparation dependencies is predictive of theta band fluctuation. These include the right and left Angular Gyrus and Amygdala, as well as the right Insula, Pallidum, Putamen, Inferior Parietal Gyrus (excl. Supramarginal and Angular Gyri), Superior Temporal Gyrus, and Middle Temporal Gyrus. The involvement of parts within the right Inferior Parietal Gyrus that are not the Angular or Supramarginal Gyrus could be due to imprecision in source localisation, with the model attributing variance to these spatially close areas that could also originate from the Supramarginal or Angular Gyri. Alternatively, the activity might be related to higher order associated processing in the right Inferior Parietal Gyrus (Obert et al., 2014). Notably, in addition to some of the areas relevant to language processing that were also previously observed in prolongation, such as the Angular Gyrus and Superior Temporal Gyrus, preparation also carried great predictive value for the Striatum. The Putamen and Pallidum are both associated with motor coordination and the reward system. This finding closely mirrors the plethora of prior studies pointing towards the involvement of the reward system in the phenomena associated with music-induced expectation, for example when an anticipated harmony is presented (Gebauer, Kringelbach, & Vuust, 2012; Huron, 2001-2002, 2006; Salimpoor, Benovoy, Larcher, Dagher, & Zatorre, 2011; Vander Elst, Vuust, & Kringelbach, 2021; Vuust & Frith, 2008). In many of these studies, the anticipation of an expected harmony is induced through chord progressions, a phenomenon that the present theoretical framework would characterise as opening a preparation dependency (when, e.g., preparatory dominants towards a final harmony are introduced). The present finding then extends this observation by showing that not only during the resolution, but also during the anticipation stage, we can use music theoretical models to make predictions about brain activity. Specifically, theta band activity in both Putamen and Pallidum decreases as the number of open preparation dependencies increases. Vice versa, this means that theta band activity increases when preparation dependencies are closed, which results in the phenomenon of dependency resolution that other studies have explored (Koelsch, Rohrmeier, et al., 2013). Future studies could reanalyse the published data, to explore whether such increases in theta band activity can indeed be observed in the Striatum when a dominant is resolved to a tonic.

In addition to the reward system, preparation also carried predictive value for areas associated with emotion processing, such as the Insula and the Amygdala. This also closely matched prior published literature. Specifically, the temporal dynamics of expectation, including its creation, its resolution and the delay of its resolution, is widely associated with the emergence of meaning and emotion in music, both from a theoretical (Meyer, 1956; Schlenker, 2017) as well as a psychological perspective (Egermann, Pearce, Wiggins, & McAdams, 2013; Huron, 2006; Sauvé, Sayed, Dean, & Pearce, 2018). Indeed, a large body of literature has been dedicated to exploring the strong emotional effects of music and much of this research highlights emotional processing areas that are active during music listening (Gosselin, Peretz, Johnsen, & Adolphs, 2007; Koelsch, 2014; Koelsch, Fritz, & Schlaug, 2008; Koelsch, Skouras, et al., 2013; Lehne, Rohrmeier, & Koelsch, 2014), in particular when prior expectations are fulfilled or denied (Cheung et al., 2019; Koelsch et al., 2008; Steinbeis, Koelsch, & Sloboda, 2006).

The involvement of the Middle Temporal Gyrus is of particular interest. This is because the left Middle Temporal Gyrus contains the auditory association cortex (Ohnishi et al., 2001) that has been found to be involved in tasks such as linking images of musical instruments to sound (Hoenig et al., 2011), attributing the correct ending to musical phrases (Groussard et al., 2010), or generating new endings to melodies (Brown, Martinez, & Parsons, 2006). The last task in particular is also associated with activity in the right Middle Temporal Gyrus, similarly to the effects observed here. As mentioned above, this tendential right-lateralisation of music processing compared to the tendential left-lateralisation of language has been observed before (Brown et al., 2006). It is possible that, when listeners are presented with an open preparation dependency, they simulate prototypical necessary completions within the syntactic rules of the idiom. This would also be consistent with psycholinguistic accounts of sentence comprehension, whereby working-memory resources are recruited when comprehenders anticipate required but not-yet-observed nodes in an online parse (Gibson, 1998; Shain, Blank, Fedorenko, Gibson, & Schuler, 2022; Shain, Van Schijndel, Futrell, Gibson, & Schuler, 2016). Since the right Middle Temporal Gyrus has been shown to be involved in generating mental simulations of future musical events (Brown et al., 2006), it may also be activated by preparation dependencies. This perspective also offers and explanation for why theta band activity in the right Middle Temporal Gyrus can be predicted with open preparation dependencies, but not prolongation dependencies. In fact, open preparation dependencies elicit an expectation towards a specific different harmony, and listeners may need to generate a mental representation of this target harmony in order to retain the prediction. Open prolongation dependencies, on the other hand, could make do with simply storing the recently encountered harmony in memory, and do not require the simulation of a new harmony.

Interestingly, in those cases where preparation and prolongation predict theta band activity for the same brain regions, the directionality of the effects is opposite. This further supports that these two types of open dependencies are processed differently. To further interpret this difference in directionality, it is important to highlight that the negative directionality in preparation specifically describes that an increase in the count of open preparation dependencies predicts lower theta band activity during the musical section that follows, compared to a decrease in open preparation dependencies which predicts an increase in theta band activity. At first glance, this may appear counter intuitive when understanding higher theta band activity as a marker of processing demand. However, this result is compatible with prior linguistic research suggesting that the process of integrating the head of an open syntactic dependency into an existing incremental representation is particularly resource intense (Gibson, 1991, 2000). It is possible that the effect we observe here is that a high integration cost is required when open preparation dependencies are closed, which increases the more simultaneously open preparation dependencies exist. This begs the question why prolongations do not contain such high integration costs, but only retention costs. This could potentially be explained by the difference in nature between these two dependencies.

Literature in psycholinguistics hypothesises that (at least) two types of cognitive operations are required to implement syntactic processing (Gibson, 1998, 2000): retaining representations of past or future events in memory (memory storage) and integrating the currently-parsed event into an existing partial parse (structural integration). Extending this framework to the musical domain, the present results are compatible with an account of musical processing whereby (1) prolongations are predominantly associated with memory storage demands extending from the time the prolonged harmony is first encountered to the time the prolongation is completed by a return to the same harmony or a substitute thereof; (2) preparations, in turn, are rather predominantly associated with the structural integration of the events that resolve the dependency, so that greater activation is predicted as the number of open preparations decreases. Memory requirements associated with preparation dependencies are qualitatively different from those associated with prolongation dependencies, as the former – and not the latter – entail encoding a projection of a future event rather than a trace of a past event. As a consequence, memory storage operations associated with preparation dependencies may recruit different brain regions, such as the right Middle Temporal Gyrus as discussed above.

### Musical Expertise

The Superior Temporal Gyrus, Angular Grus, and Heschl’s Gyrus all showed a moderating effect of musical expertise. We believe three observations are important to highlight. First, the observed mediating effects of musical expertise were exclusively left lateralised. Second, the observed effects were nearly exclusively related to prolongation. Third, the effect directionality suggests stronger effects in non-Musicians compared to Musicians. Taken together, we believe the most compelling explanation is that the observed musical expertise effect is predominantly driven by general memory processes. Prior studies have repeatedly shown that musical expertise can affect memory for musical as well as non-musical stimuli with musicians generally outperforming non-musicians (Franklin et al., 2008; George & Coch, 2011; Hansen, Wallentin, & Vuust, 2013; Oechslin, Van De Ville, Lazeyras, Hauert, & James, 2013), and often recruiting less resources whilst doing so (Fujioka, Trainor, Ross, Kakigi, & Pantev, 2004; Herff & Czernochowski, 2019; Moradzadeh, Blumenthal, & Wiseheart, 2015). Consequently, the additional memory costs of open prolongation dependencies likely disproportionally burden non-musicians, further strengthening the predictive value of the number of open prolongations for theta activity. Since this explanation assumes that non-musicians also engage with tracking harmonic dependencies, this would suggest that at least some aspects of musical syntactic processing in the domain of harmony are automatic (Justus & Bharucha, 2001) and rely on implicit rather than explicit knowledge (Reber, 1989). Rather than influencing whether a parsing process is attempted in the first place, musical training may then manifest itself in how successful and how effortless it is. This observation is of particular interest, as due to the monophonic surface of the stimuli, listeners had to infer harmonic function, suggesting that latent harmonic entities and relationships among them are indeed a reasonable abstraction of the musical surface even for monophony.

However, the categorical classification of musicianship used in this study lacks nuance. Future studies should evaluate the effect of musical expertise on brain activity involved in the structural processing of prolongation and preparation in greater detail, for example by deploying more nuanced measures of musical expertise and comparing how linguistic expertise affects brain activity during parsing of linguistic structural with the effect of musical expertise on the parsing of structural dependencies in music.

### Limitations and Future directions

We split the analysis into two parts, pre- and post-harmonic rhythm change. This was done due to computational power limitations, as well as to account for potential differences in the difficulty for listeners to infer the underlying harmonies in the two parts. However, since variable task difficulty was not a research question in this study, and instead we were concerned with providing a general proof-of-concept and identifying regions of interest, we interpreted the combined results. That been said, there are clearly some differences between the two parts of the piece, including a rather puzzling flip in the sign of the effect observed in the Pallidum. There is a plethora of striking musical differences between these two parts of the piece, and any of them as well as their combination might justify different responses in terms of affect or motor control. Additional qualitative reflection on the music-theoretical analysis of the piece may further result in better understanding why, in some parts of the piece, clearer results were obtained for prolongation relative to preparation or vice versa. However, such analysis is beyond the scope of this article and would likely involve, on one hand, traditional music-theoretical approaches to engage with and complement the results of the present study and, on the other hand, a new data collection as well as a sophisticated quantification of perceptual harmonic clarity, i.e., how clearly the monophonic musical surface as presented is segmented into an abstract symbolic representation in terms of chords.

Despite much prior evidence of the involvement of Broca’s area in syntactic processing of music (Maess et al., 2001), we did not observe additional evidence that prolongation and preparation carry predictive value for theta-band activity in Broca’s area. We suspect that the explanation lies in the fact that prior studies predominantly utilised paradigms based on instantaneous syntactic outliers or violations (Koelsch, Gunter, Wittfoth, & Sammler, 2005; Koelsch, Jentschke, Sammler, & Mietchen, 2007; Koelsch, Maess, & Friederici, 2000; Maess et al., 2001; Sammler, Koelsch, & Friederici, 2011), rather than continuously varying embedding depth of syntactically valid musical phrases. As a consequence, it may be argued that the role of Broca’s area is tied to maintaining and evaluating an active structural representation (Koelsch, 2006), responding to parsing difficulty or failure when necessary (Patel, Gibson, Ratner, Besson, & Holcomb, 1998) and potentially triggering processes of syntactic repair or revision in cases of mismatch (Cecchetti, Herff, & Rohrmeier, 2022), rather than to the computations pertaining to incremental syntactic integration. This is also consistent with evidence from language and music that the specific role of Broca’s area during online parsing manifests itself as a form of cognitive control (Slevc & Novick, 2013). It is important to note that whilst the present finding suggest that a representation of syntactical musical structure exists, at least in terms of correlates of embedding depth and dependency types, this representation is not necessarily encoded explicitly but could rather be implicit, with varying degree of conscious awareness.

Combining the literature reviewed throughout this paper with the present results paints a picture of the processing of syntactic structure in music that can be summarised as follows.

A network of areas containing the Angular Gyrus, Heschl’s Gyrus, Supramarginal Gyrus, and Superior Temporal Gyrus are involved in the segmentation and analysis of the hierarchical syntactic musical structure, whereas the Broca’s area plays a key role in the evaluation of grammaticality. Regions such as the Amygdala, Insula, and Striatum are then involved in the processing of the emotional and reward responses evoked by the syntactic structure, such as fulfilled or denied expectations. Some of these areas are shared between music and language, such as the left Angular Gyrus, Heschl’s Gryus, and Supramarginal Gyrus, whereas others are distinct, such as a number of areas in the right hemisphere that correspond to language-processing areas in the left hemisphere. The precise areas that are recruited depends on the type of open dependency (prolongation vs. preparation), with prolongation relying on many established auditory (Heschl’s Gyrus) and language-related areas (e.g., Angular Gyrus and Supramarginal Gyrus) while preparations rely more on areas involved in generating precise predictions towards non-identical events (e.g., right Middle Temporal Gyrus) and emotional processing of the fulfilment or denial of these expectations (e.g., Insula, Pallidum, Putamen). Features of the musical surface (e.g., audio envelope) as well as musical expression (e.g., differences between renditions of the same piece) then likely stand in a reciprocal relationship with musical structure, with all these parameters further mediating the effects of each other.

## Conclusion

The present investigation extents to the musical domain prior linguistic research showing that the processing of syntactic embedding is reflected in brain theta power. Whilst various brain areas among those identified in this study are comparable to those observed in linguistic research, others are different or lateralised in the right instead of the left hemisphere. In addition to extending previous linguistic findings to the musical domain, this study also contains some additional contributions. Firstly, prior studies predominantly focused on comparing embedded and non-embedded conditions, whereas here we continuously model the precise depths of embedding, as quantified within a music-theoretically motivated framework. We then show that embedding depths also predicts theta activity continuously and before the points of resolution We show that the accumulation of open dependencies indeed continuously influences processing as the number of open dependencies increases. Secondly, due to the precise music theoretical framework, we can differentiate between different types of embedding and show that they are associated with different effects in only partially overlapping brain areas. Thirdly, we show evidence for the process of harmonic dependencies even harmony has to be inferred from a monophonic surface. Taken together, and similarly to prior studies in linguistic, the present study functions as a proof-of-concept showing that precisely formalised music theoretical models can be used to predict brain theta power in listeners, suggesting that a representation of the musical structure, akin to that described by the music theoretical model, is formed during listening to a specific, familiar musical idiom.

## Author contributions

SH, GC, MR conceived of the concept, the study, and the methods. LB, PV, LMK provided the experimental data. LB conducted pre-processing and the source localisation of the MEG data and contributed to the visualizations. SH implemented the model and conducted data analysis. GC and MR analysed the piece and computed the music theoretical predictors. SH wrote the first version of the manuscript and revised it. GC and LB contributed to the writing. GC, MR revised the manuscript; PV and LMK commented on the manuscript.

## Acknowledgments

This project has received funding from the European Research Council (ERC) under the European innovation program under grant agreement No 760081-PMSB and by Mr Claude Latour through the Latour Chair of Digital and Cognitive Musicology. SAH is supported by the Australian Government through the Australian Research Council (ARC) under the Discovery Early Career Researcher Award (DECRA, DE220100961) The Center for Music in the Brain (MIB) is funded by the Danish National Research Foundation (project number DNRF117). LB is supported by Carlsberg Foundation (CF20-0239), Lundbeck Foundation (Talent Prize 2022), Center for Music in the Brain, Linacre College of the University of Oxford and Nordic Mensa Fund. MLK is supported by Center for Music in the Brain and Centre for Eudaimonia and Human Flourishing, which is funded by the Pettit and Carlsberg Foundations.

